# Development of a *Xenopus*-based assay for high-throughput evaluation of mucociliary flow

**DOI:** 10.1101/2022.07.12.499724

**Authors:** Ioanna Antoniades, Andria Koulle, Maria Chatzifrangkeskou, Timothea Konstantinou, Paris Skourides

## Abstract

Motile cilia are organelles lining the surfaces of major organs of the human body and generate directional fluid flow. Ciliary dysfunction has been linked to an emerging class of multisystem disorders, collectively known as motile ciliopathies. Drug screening for ciliopathies is challenging due to the unavailability of high-throughput assays that can evaluate ciliary flow generation. Here, we describe the development of a unique assay that enables the direct and rapid evaluation of mucociliary flow, which simultaneously facilitates high-throughput screening of potential therapeutic agents for motile ciliopathies. The assay relies on the ability of *Xenopus* tadpoles to promote mixing of a two-phase differential density aqueous mixture, through the robust flow generated by the mucociliary epithelium on their epidermis. We show that the rate of phase mixing is proportional to the rate of cilia-driven flow, therefore it directly represents the effectiveness of flow generation. We also demonstrate that the assay can detect changes in ciliary flow elicited by defects in cilia, CBF modulation and rotational polarity, providing an ideal assay for the identification of CBF-modulating compounds, as potential drugs for motile ciliopathies. Importantly we use the assay to show that CBF modulating drugs can improve flow generation and could thus be used as a potential therapeutic approach in PCD patients. The assay we have developed thus represents a powerful new tool for research, as well as drug development.

## Introduction

Mucociliary clearance constitutes the primary innate defense mechanism of the airways and is primarily driven by multiciliated cells (MCCs). MCCs possess hundreds of motile cilia that project outwards the apical surface of cells and beat in metachronal waves, thus generating directed flow that drives inhaled particles and pathogens out of the respiratory tract (Bustamante-Marin and Ostrowski, 2017). Outside the airways, motile cilia line major organs of the human body, including the brain ventricles and the oviducts, where they mediate circulation of the cerebrospinal fluid and gamete and embryo transport, respectively (Ibanez-Tallon et al., 2003; Roy, 2009). Generation of directional flow is regulated by multiple parameters, including proper structure and composition of cilia, that influence the pattern and frequency of beating, as well as proper alignment of cilia, that governs the polarity and directionality of flow(Bush et al., 1998; Marshall and Kintner, 2008; Meeks and Bush, 2000; Noone et al., 2004).

Dysfunction of motile cilia is linked to a variety of disorders in humans, referred to as motile ciliopathies, leading to recurrent respiratory tract infections, hydrocephaly and infertility (Noone et al., 2004). Primary ciliary dyskinesia (PCD) is the most well studied multisystem disorder, characterized by impaired mucociliary clearance. PCD is genetically and phenotypically heterogeneous, and patients usually suffer from chronic lung infections, sinusitis, bronchiectasis and subfertility (Ibanez-Tallon et al., 2003; Noone et al., 2004). In most of the patients, dysfunction stems from defects in the ultra-structure of motile cilia, such as defects in the dynein arms and the central axonemal microtubules, that affect cilia motility and therefore mucociliary clearance (Ibanez-Tallon et al., 2003; Noone et al., 2004). Interestingly, abnormal motility is sometimes accompanied by disorientation of cilia, which however is primarily considered to be a secondary effect or the consequence of defective flow generation (Jorissen and Willems, 2004; Marshall and Kintner, 2008; Rautiainen et al., 1990).Proper ciliary orientation is a fundamental property of multiciliated tissues, ensuring the unidirectionality of flow. This is achieved by the uniform orientation of cilia both within single cells (rotational polarity) and maintained throughout the mucociliary epithelium (tissue polarity). While cues from the planar cell polarity (PCP) pathway and cytoskeletal dynamics have been shown to govern the initial establishment of polarity, hydrodynamic forces and a positive feedback mechanism, in response to flow, reinforce the orientation, further refining polarity along the tissue (Marshall and Kintner, 2008; Mitchell et al., 2009; Vladar et al., 2012). Therefore, acquired disorientation of cilia is typical in patients with motility defects (Jorissen and Willems, 2004). However, other studies showed cilia misorientation and inefficient mucociliary clearance, in individuals with otherwise normal ciliary beating and structure, suggesting that orientation abnormalities may represent a sub-class of PCD that remains to be explored (Marshall and Kintner, 2008; Rayner et al., 1996).

The genetic and phenotypic heterogeneity of PCD make disease diagnosis and management challenging. Most therapeutic approaches have focused on symptomatic relief by reducing sputum density and providing early treatment of infections. A number of airway clearance techniques, combined with inhaled bronchodilators and agents reducing mucus density, are used to promote cough and therefore mucus removal. On the other hand, antibiotic and anti-inflammatory treatments are used to manage infections (Bryson and Sorkin, 1994; Lucas et al., 2014; Rubbo and Lucas, 2017). However, early initiation of symptomatic treatment has not yet resulted in an unequivocal stabilization or improvement of the disease course. Compounds that increase the CBF are expected to improve mucociliary flow (Braiman and Priel, 2008); hence, future treatment with drugs that target cilia directly, instead of mucus consistency, could have a beneficial effect on mucociliary clearance, and ultimately disease progression in these patients. Importantly, a number of compounds have been identified as ciliostimulatory agents, that increase CBF in mammalian cells in culture, however their impact on mucociliary clearance itself has not been assessed (Grosse-Onnebrink et al., 2016; Haxel et al., 2001; Joskova et al., 2020; Workman and Cohen, 2014; Yasuda et al., 2020).

The internal positioning of multiciliated epithelia in mammals prohibits the functional evaluation of the effectiveness of CBF promoting compounds. Therefore, assessment of the effectiveness of flow primarily relies on human cells cultured in air liquid interface (ALI) systems (Workman and Cohen, 2014). Such methods focus on calculating local flow velocity, by tracking individual fluorescent particles placed in the culture medium, over time. Even though a number of factors affecting flow speed (such as CBF and CBP) can be measured in ALI culture systems, other parameters (such as tissue-wide cilia orientation) are impossible to detect. This stems from the fact that due to the lack of tissue polarity-guiding cues, MCCs are randomly polarized with respect to each other, therefore restricting live observations to isolated cells or very small areas of the tissue.

Similar to mammalian airway mucociliary epithelia, the epidermis of *Xenopus* tadpoles consists of a polarized mucociliary epithelium, which resembles that of the mammalian respiratory tract. *Xenopus* MCCs generate directional flow over the skin. Effective mucociliary clearance is critical since the tadpoles respire through their skin and cilia driven flow is important for pushing away dirty mucus and waste products as well as for refreshing oxygenated water over the epidermis (Steinman, 1968; Werner and Mitchell, 2012). In the past decades a plethora of studies have demonstrated the advantages of *Xenopus* as a model for the study of MCC differentiation and function. The external nature of the ciliated epithelium in *Xenopus*, uniquely enables the study of ciliogenesis and ciliary function *in vivo* and facilitates live observation and measurement of mucociliary flow at the tissue level (Werner and Mitchell, 2012). This is achieved by high-speed video recording of the displacement of fluorescent particles placed in the tadpole medium, followed by computational tracking of individual beads, determination of flow polarity and calculation of flow speed. Although powerful, this approach is labor intensive and cannot be applied in a high-throughput scale, to screen chemical compound libraries for potential PCD drug candidates.

In this study we took advantage of the robust flow generated by the epidermal MCCs of *Xenopus* tadpoles and developed a novel assay that enables automated quantitative evaluation of mucociliary flow in a fast, accurate and reproducible manner. We show that the flow generated by tadpoles promotes the mixing of a biphasic system. The system comprises of a fluorophore and a quencher, each incorporated in one out of two phases of differential density. Flow-driven mixing of the two phases promotes the interaction between the fluorophore and the quencher. Mixing is directly measured as a decrease of fluorescence intensity. We go on to show that the rate of mixing is proportional to the flow velocity and that the assay can detect and quantify changes in velocity, resulting from defective mucociliary function. In addition, we provide evidence that the assay has the capacity to track changes in parameters that influence mucociliary flow in different ways. Specifically, we show that it can detect changes in CBF that quantitatively impact flow velocity, while at the same time it can identify cilia polarity defects, which qualitatively affect flow. Finally, we demonstrate that the *Xenopus*-based flow evaluation assay has the capacity to be applied for multiscale screening studies and validate the effects of potential drug candidates for PCD.

## Materials and Methods

### Embryo Obtainment and Manipulation

Female adult *Xenopus laevis* frogs were ovulated by injection of 750U human chorionic gonadotropin. Eggs were fertilized *in vitro* and de-jellied in 2% cysteine (pH 7.8). Embryos were maintained in 0.1× Marc’s modified ringers (MMR) and staged according to Nieuwkoop and Faber (P.D. Nieuwkoop, 1994).

All procedures were carried out according to the University of Cyprus Animal Ethics Guidelines under licenses granted by the Animal Health and Welfare Division of the Veterinary Services department of the Ministry of Agriculture, Rural Development, and Environment of the Republic of Cyprus.

### FAK morpholino (MO) microinjections

Microinjections were performed in 4% Ficoll in 0.33× MMR using glass capillary pulled needles, forceps, a Singer Instruments MK1 micromanipulator and a Harvard Apparatus pressure injector. Injected embryos were reared for 2h or until stage 8 in 4% Ficoll and then washed and maintained in 0.1 × MMR. 7.5ng or 15ng FAK MO were injected at each ventral blastomere of 4-cell stage embryos (at the animal pole to target the epidermis) together with 75pg centrin4 RFP, which was used as a linage tracer to evaluate the uniformity of MO injections. Embryos were allowed to develop to the appropriate stage and then fixed in MEMFA (Hazel L. Sive, 2000) for 2h at room temperature or were used to assess mucociliary flow.

The sequence of FAK MO used in our experiments is TTGGGTCCAGGTAAGCCGCAGCCA (Fonar et al., 2011).

### Whole mount Immunofluorescence

Immunofluorescence experiments were performed as previously described (Antoniades et al., 2017; Antoniades et al., 2014). Briefly, fixed embryos were permeabilized in PBDT (0.5% Triton X-100, 1% DMSO in 1× PBS) and blocked in 10% normal donkey serum for 1h at room temperature. Primary antibody was added and incubated for 4h at room temperature. The primary antibody used was acetylated α-tubulin (6–11B-1, Santacruz Biotechnology). Embryos were washed in PBDT and incubated in secondary antibody (Alexa 488, Invitrogen) for 2h, at room temperature. Stained embryos were washed in PBDT, post-fixed in MEMFA for 20min at room temperature, washed in 1× PBS, and imaged immediately on a Zeiss LSM 710 laser scanning confocal microscope with the Zen 2010 software.

### Drug treatments

Embryos were treated with 10μM Cytochalasin D or 1μM Nocodazole between stages 23 and 29, at 16°C, as previously described (Werner et al., 2011). Control embryos were treated with 0.1% DMSO.

### CBF-modulating compound treatments

Stage 34 embryos were treated in 20mMcAMP, 10ng/ml IL-5, 30ng/ml IFN-_γ_, or 80 μM Prostaglandin-E2 for 1h, at room temperature.

### Flow velocity assessment and quantification

Stage-34 tadpoles were anesthetized in 0.01% benzocaine in 0.01× MMR and placed in custom-made PDMS wells on glass slides. QDs (QDot 655, Invitrogen) were added into MMR and a glass coverslip was placed on top of the wells. The wells were thick enough so as to avoid contact between the coverslip and the skin of the tadpole. Imaging of QD displacement was performed on a Zeiss Axio Imager Z1 microscope equipped with a Zeiss Axiocam MR3 and the Axiovision software 4.7, using the Fast Acquisition mode. The IMARIS software was used to automatically track the displacement of individual QDs and quantify the flow velocity.

### CBF analysis

Image of cilia beating was performed on a ZEISS Axiovert 200M microscope equipped with a Basler scA640 video camera. Analysis and measurement of CBF were performed using the Sisson-Ammons Video Analysis System.

### Evaluation of Xenopus-driven mixing of solutions of differential density

Stage 34 embryos were anesthetized in 0.01% benzocaine in 0.01 × MMR for 5min. Embryos were transferred into individual wells of a 96-well plate and excess MMR was removed. 100 μl of 50 nM Biotin-4-FITC (Santa Cruz, sc-319858), 0.01% benzocaine in 0.1 x MMR was introduced into each well (low-density solution). 6μl 12,5nM streptavidin (Invitrogen, #434301) in 25% Ficoll (Cytiva #17030050) (in 0.1 x MMR) were carefully introduced into the bottom of each well (high-density phase). 3-5 wells containing the differential density solutions without a tadpole, were included in each set of experiments as negative controls. 3-5 wells in which the differential density solutions were mechanically mixed with a micropipette, were included in each set of experiments as positive controls.

For real-time monitoring of fluorescence intensity, plates were immediately transferred into a TECAN infinite f200 plate reader equipped with a Exc485/Emm510 filter and fluorescence was measured every 5 min for 1h.

For end-point measurement of fluorescence intensity, the embryos were incubated in the biphasic solution for 30 min and 100 μl of 500 nM D-Biotin (Sigma-Aldrich) were added and mechanically mixed with the solution. Fluorescence was measured using a TECAN infinite f200 plate reader.

For quantification, all measurements were normalized in order to show the percentage of fluorescence change over time.

### Statistical analysis

Graph generation and statistical analysis were performed using the Prism software (GraphPad, San Diego, CA). Graph data are presented as mean values and error bars represent S.E.M. Statistical analysis was performed using two-tailed unpaired t-tests, with 95% confidence interval.

## Results

### The robust flow generated by the Xenopus mucociliary epithelium promotes the mixing of solutions with different density

Current methods of assessing mucociliary flow are time consuming, and labor intensive. As such they cannot be applied to libraries to screen for potential drug candidates. In order to provide a quick and easy evaluation of mucociliary function, we investigated the possibility that cilia-driven flow generated by *Xenopus* tadpoles promotes liquid mixing of a biphasic solution. Specifically, we postulated that robust flow, generated by the mucociliary epithelium comprising the epidermis of *Xenopus* tadpoles, would speed up the mixing of a system composed of two phases of differential density. The system consists of a low-density phase, composed of 0.01% benzocaine in 0.1 x MMR and a high-density phase, composed of 25% Ficoll in 0.1 x MMR with 0.01% benzocaine. Ficoll was selected because of its high molecular weight and low content of dialyzable material, giving solutions of low osmolality and osmotic pressure, as well as its low cell membrane permeability. The concentration of Ficoll required to prepare the high-density solution was experimentally determined. Specifically, a series of solutions of different concentrations were examined for their ability to generate a visually distinct high-density phase, which exhibited minimal mixing with MMR over time. In order to facilitate visualization of mixing, we introduced Quantum Dots to label the high-density (bottom) phase (**Figure 1**).

**Figure 1:**
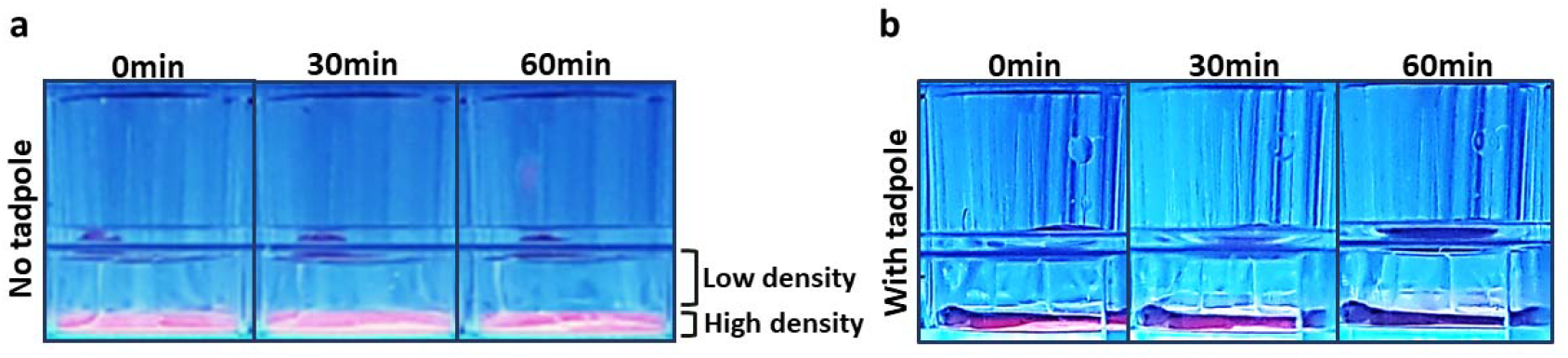
Xenopus cilia driven flow promotes the mixing of solutions of differential density. Images of plastic wells containing two solutions of differential density; a high-density solution composed of 25% Ficoll, in 0.1 x MMR and fluorescently labeled with QDs, and a low-density solution composed of 0.1 x MMR, with (b) or without a tadpole (a). Because of their differential density the solutions create visually discrete phases. In the absence of a tadpole, no significant mixing of the two solutions is observed over time (a). *Xenopus* tadpoles promote gradual mixing of the two solutions, as the two phases become less discrete over time (b).

The low-density solution was initially added into the wells of a 96 well-plate, followed by the introduction of *Xenopus* tadpoles (stage 34) into individual wells. The high-density solution was then carefully introduced to the bottom of the wells. Wells devoid of any tadpole were used as controls and represent the diffusion-driven mixing. Images of the wells were acquired every 5 minutes, using a UV trans-illuminator, for a total of 1 hour. As shown in **Figure 1**, unlike control wells where nearly no mixing is observed, in the presence of a tadpole the two phases were gradually mixed. This is evident by the diffusion of the QD fluorescence signal and the generation of a weak fluorescent interphase over the tadpole, after the 1-hour incubation. This shows that the robust flow generated by the *Xenopus* mucociliary epithelium can promote the mixing of solutions with different densities and could thus be used as a basis for flow quantification.

### FITC-fluorescence quenching enables easy readout and automated end-point quantification of phase mixing

Although enabling the detection of flow generation, the set-up described above does not allow the quantification of the extent of mixing. Therefore, it cannot be used to measure flow effectiveness, preventing its application for the detection of minor changes in flow rate in a liable manner. In order to facilitate easy and accurate readout of mixing that can be adapted for high-throughput applications, we took advantage of the Biotin-4-FITC/Streptavidin quenching system. Because of its small size, Biotin-4-FITC shows exceptionally fast and tight binding towards streptavidin, accompanied by strong quenching and measurable loss of FITC fluorescence(Kada et al., 1999a; Kada et al., 1999b). The system has been used to measure avidin and streptavidin concentrations in crude biofluids, to determine the biotin-binding capacity of streptavidin conjugated QDs, to characterize neutravidin-modified nanoparticles as drug carrier systems and to determine D-biotin doses in medical treatments, by quantifying the extent of fluorescence quenching (Balthasar et al., 2005; Ebner et al., 2008; Gruber et al., 1998; Kada et al., 1999b; Mittal and Bruchez, 2011). We therefore postulated that introduction of streptavidin and Biotin-4-fluorescein-FITC into the differential-density solutions will be followed by quenching only upon mixing, which is expected to promote Biotin-4-FITC binding to streptavidin (**Figure 2a**). Quantification of fluorescence intensity changes would be expected to correlate to the extent of mixing and therefore the effectiveness of mucociliary flow.

**Figure 2:**
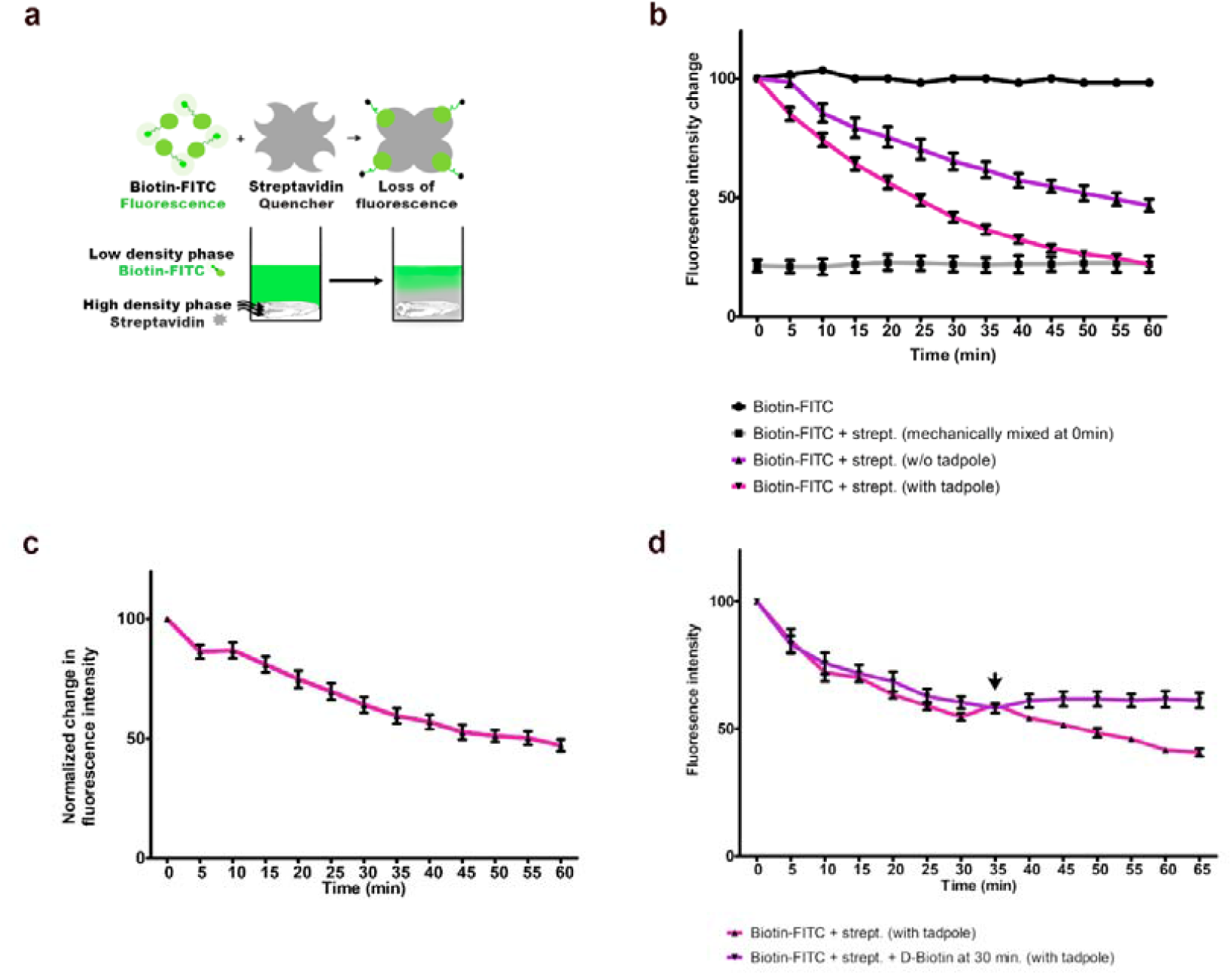
FITC-fluorescence quenching enableseasy readout and automated quantification of mixing efficiency. a) Introduction of Biotin-4-FITC in the low-density solution and streptavidin, acting as a quencher, in the high-density solution, enables automatic quantification of changes in fluorescence, as a result of mixing of the two solutions followed by FITC quenching. b) Fluorescence intensity change in the presence (pink chart, n=7) or the absence (purple chart, n=10) of a *Xenopus* tadpole. Cilia-driven flow from *Xenopus* tadpoles promotes faster mixing and therefore fluorescence quenching, that diffusion. Mechanical mixing of the biphasic system (resulting in drastic quenching) is used as positive control and Biotin-FITC without streptavidin is used as negative control. c) Quantification of the ratio of fluorescence intensity in the presence of a tadpole, to fluorescence intensity in the absence of a tadpole, reveals fluorescence changes resulting exclusively from flow-driven mixing of FITC (low-density) and streptavidin (high-density) solutions. d) Normalized change in fluorescence intensity as described in 2c. Addition of excess biotin (indicated by the arrow) and mechanical mixing of the solution, at 30 minutes (purple chart, n=10), stops FITC-biotin/streptavidin interaction and therefore prevents further FITC quenching. This facilitates endpoint, automatic measurement of fluorescence intensity, enabling high-throughput application of the assay.

In order to investigate this possibility, we prepared 0.1 x MMR and 25% Ficoll in 0.1 x MMR solutions containing Biotin-4-FITC and streptavidin, respectively. The amount of Biotin-4-FITC and streptavidin used were experimentally determined in order to ensure that maximum sensitivity and dynamic range for the detection of changes in flow is achieved. Specifically, the optimal amount of FITC was selected so as to display at least 10 times higher intensity than the minimum detectable signal by an automatic plate reader, whereas the amount of streptavidin was selected based on its potential to allow minimum fluorescence change in the absence of mixing, while achieving maximum quenching upon mechanical mixing with the low-density solution. *Xenopus* tadpoles (stage 34) were immersed in the Biotin-4-FITC solution (low density-phase) in individual wells of a 96-well plate. The streptavidin solution (high-density phase) was introduced at the bottom of each well, generating a bi-phasic Biotin-4-FITC/streptavidin system. Negative (Biotin-4-FITC only) and a positive (Biotin-4-FITC + streptavidin, mechanically mixed with a micropipette) control wells were included in each plate, for each experiment. In order to account for any non-cilia mediated mixing of the solutions, or Brownian motion-mediated Biotin-4-FITC/streptavidin binding, an additional negative control well, containing the biphasic system in the absence of a tadpole, included in each plate. The plates were immediately introduced into an automatic plate reader and fluorescence intensity in each well was measured every 5 minutes for a total of one hour. As shown in **Figure 2b**, a slight decrease of FITC intensity is observed in empty wells suggesting a passive mixing of the two phases and transition of the two molecules between the phases. However, *Xenopus* tadpoles and therefore mucociliary flow, drastically speed-up the mixing, as indicated by the higher rate of fluorescence drop in tadpole-containing wells, compared to the ones without. Importantly, maximum quenching is achieved in the presence of a tadpole, after one-hour incubation (**Figure 2b**). In order to account for the non-flow driven fluorescence change, we normalized our data by calculating the ratio of flow-driven (with tadpole) to non-flow driven (without tadpole) fluorescence intensity, for each time point. Quantification of the normalized changes in fluorescence intensity (**Figure 2c**) suggests that the robust cilia-driven flow from the tadpoles accelerates mixing, facilitating easy mixing-readout as measurable FITC quenching.

In order for this approach to be applied in a quick and high-throughput scale though, it should ideally enable simultaneous quantification of multiple samples. Our data (**Figures 2b and c**) show that the extent of mixing is proportional to incubation time, requiring continuous real-time monitoring of fluorescence intensity on a plate reader. This limits the capacity of the assay to one plate at a time, since all samples of an experiment in one plate should be evaluated before a second set of samples is set. To counteract this limitation, we decided to examine the possibility of making this an end-point assay by terminating the extended Biotin-4-FITC/streptavidin interaction, at a time point where fluorescence intensity drops sufficiently but is not completely quenched, so as to ensure the discrimination of fluorescence intensity changes under different conditions. To achieve this, we used D-biotin in excess, which was introduced into the system and mechanically mixed with the solutions. Excess D-biotin instantly occupies all remaining binding sites on streptavidin, preventing further interaction with Biotin-4-FITC and thus terminating fluorescence quenching. 10-fold excess biotin was added after 30 minutes of incubation, when about 50% decrease in FITC intensity is observed in wells with tadpoles (**Figure 2b and d**). As shown in **Figure 2d**, addition of excess biotin (indicated by the arrow) prevents further decrease in fluorescence intensity which remains stable, unlike the negative control (no excess biotin) where FITC continues to quench over time. These data show that addition of excess biotin can effectively stop the Biotin-4-FITC/streptavidin interaction, enabling end-point measurement of fluorescence intensity. Using this approach, quantification of flow rates can be carried out as an end point measurement, facilitating the application of the assay in large scale experiments.

### The Xenopus flow evaluation assay can detect and quantify changes in flow generation resulting from defects in multiciliated cells

We then asked if this assay has the dynamic range and sensitivity to detect changes in flow generation stemming from defective MCC function. To investigate this possibility, we evaluated its capacity to discriminate between normal and defective flow in control tadpoles and Focal adhesion kinase (FAK) morphants, respectively. Previous work from our lab has identified FAK as a component of multiprotein complexes, named ciliary adhesions. FAK together with other ciliary adhesion proteins, associate with the basal bodies, mediating complex interactions between cilia and actin in MCCs. Downregulation of FAK disrupts basal body-actin interaction, inducing dose-dependent defects in MCC differentiation and MCC-driven flow (Antoniades et al., 2017; Antoniades et al., 2014).

Here, we used control tadpoles and tadpoles injected with either 15ng or 30ng of a well-characterized FAK MO. The effects of the MO were validated by staining and imaging cilia density and MCC morphology, as previously described (**Supplementary figure 1**) (Antoniades et al., 2017; Antoniades et al., 2014). Embryos of each group were then introduced into wells with the biphasic Biotin-4-FITC/streptavidin system, and FITC intensity in individual wells was either measured every 5 minutes, over a period of one hour (**Figure 3a**), or upon addition of excess biotin after 30 minutes of incubation (end-point measurement, **Figure 3b**). As shown in **Figure 3 a and b**, control tadpoles display higher rate of fluorescence quenching and therefore higher rate of mucociliary flow-promoted mixing, compared to FAK morphant tadpoles. Importantly, the assay can clearly distinguish different extents of flow deficiency, as evident by slower quenching rate in embryos injected with higher amounts of FAK MO (magenta chart), compared to those injected with lower amounts (grey chart) (**Figure 3 a and b**). In order to verify that the different rates of mixing resulted from impaired ciliary flow, we went on to directly measure flow speed in the different groups of tadpoles. This was achieved through high-speed video recording of fluorescent particles and quantification of the displacement rates. As shown in **Figure 3 c**, significant dose-dependent reduction in flow velocity is observed upon downregulation of FAK and this is proportional to the extent of FITC quenching, and therefore mixing efficiency, demonstrated in **Figures 3 a and b**.

**Figure 3:**
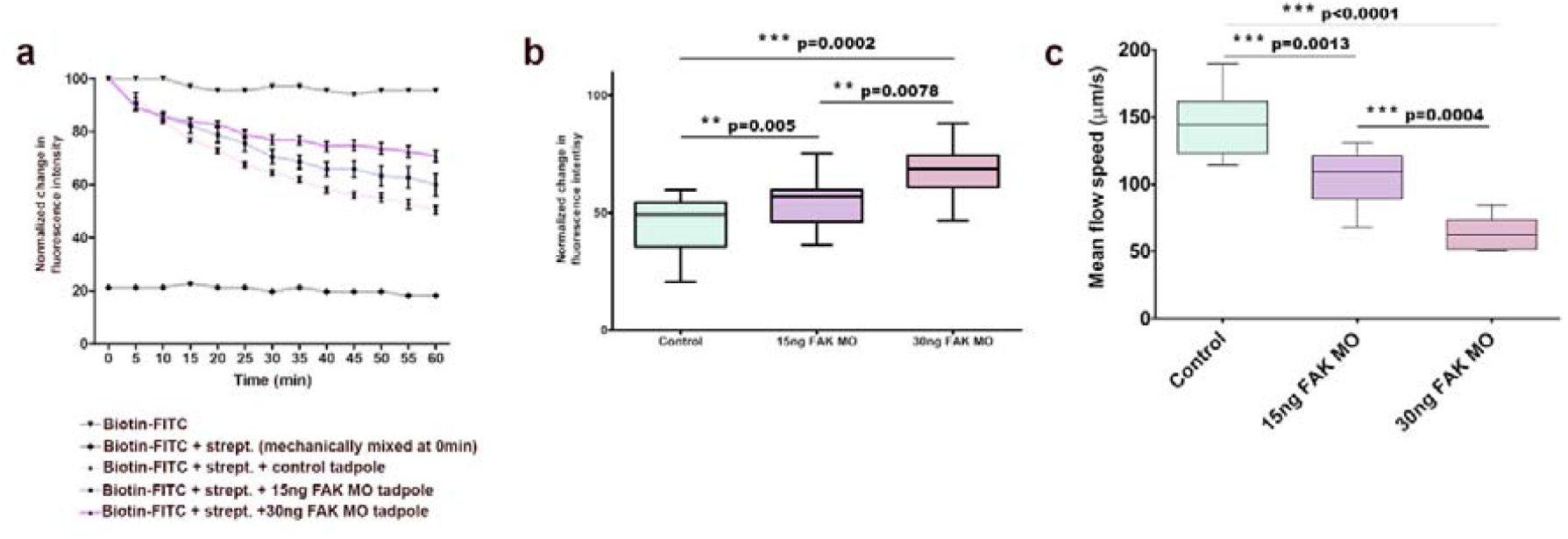
The Xenopus flow evaluation assay identifies and quantifies defects in mucociliary flow. a-b) Mixing efficiency of control tadpoles (n=12) and tadpoles injected with 15ng FAK MO (n=12), or 30ng FAK MO (n=8). a) Quantification of changes in FITC intensity over time. b) End-point quantification of FITC intensity, following addition of excess biotin at 30min of incubation. c) Quantification of flow velocity in control tadpoles (n=9) and FAK morphant embryos (15ng MO n=10, 30ng MO n=6). Downregulation of FAK in *Xenopus* epidermis leads to dose-dependent ciliogenesis defects followed by defective flow. The rate of mixing demonstrated in a and b, is proportional to the velocity of flow, shown in c.

Collectively, these results provide strong evidence that the assay we have developed provides sufficient sensitivity to track changes in flow velocity, enabling rapid and accurate quantitation of flow generation by the *Xenopus* mucociliary epithelium.

### The Xenopus flow evaluation assay provides linear responses to changes in flow

The flow evaluation assay we describe can detect changes in flow attributed to aberrant cilia formation. However, even in intact tissues with normal cilia density, additional parameters can affect the effectiveness of flow. CBF has been directly correlated to the velocity of flow, and changes in CBF have been shown to result in linear changes in flow (Christopher et al., 2014; Sears et al., 2015; Smith et al., 2011). For this reason, CBF modulating compounds have been proposed as potential treatment for PCD patients (Joskova et al., 2020). In order to assess the sensitivity of the assay to track changes in flow stemming from changes in CBF, we took advantage of the fact that CBF can be directly regulated by temperature in a linear fashion. In fact, both in human nasal epithelial and mouse tracheal cells, any temperature increase, or decrease is followed by respective enhancement or decline of CBF (Christopher et al., 2014; Salathe, 2007; Sears et al., 2015; Smith et al., 2011). To determine the impact of temperature changes on the *Xenopus* mucociliary epithelium, we evaluated i) the CBF, using brightfield high-speed video microscopy followed by frequency analysis and ii) the flow velocity, using high-speed fluorescence video microscopy to image and analyze the displacement of fluorescent particles, within a range of temperatures (14-24°C). As shown in **Figure 4 a and b**, both CBF and flow velocity display linear correlation with temperature similarly to what has been suggested for mammalian cilia. This is better represented in Figure 4 c, showing that higher CBF results in increased flow rates. We went on to examine the capacity of the *Xenopus* flow assay to detect such changes. Embryos were introduced into wells with the biphasic Biotin-4-FITC/Streptavidin system and plates were incubated at 14°C, 20°C or 25°C. After 30-minute incubation and addition of excess biotin, fluorescence intensity changes, under each condition, were determined using an automated plate reader. Each plate contained wells with tadpoles that were used to measure CBF, immediately after incubation. As shown in **Figure 4d**, increase in CBF is accompanied by faster mixing of the solutions, evident by more pronounced FITC quenching. Overall, these data show that changes in the speed of flow, resulting from CBF modification, can be sufficiently tracked and quantified using the *Xenopus* flow evaluation assay. For example, 60% increase in CBF (from 12 Hz to 19 Hz) is followed by 77% increase in flow velocity (from 152 μm/sec to 270 μm/sec), which results in 20% faster mixing of the biphasic system.

**Figure 4:**
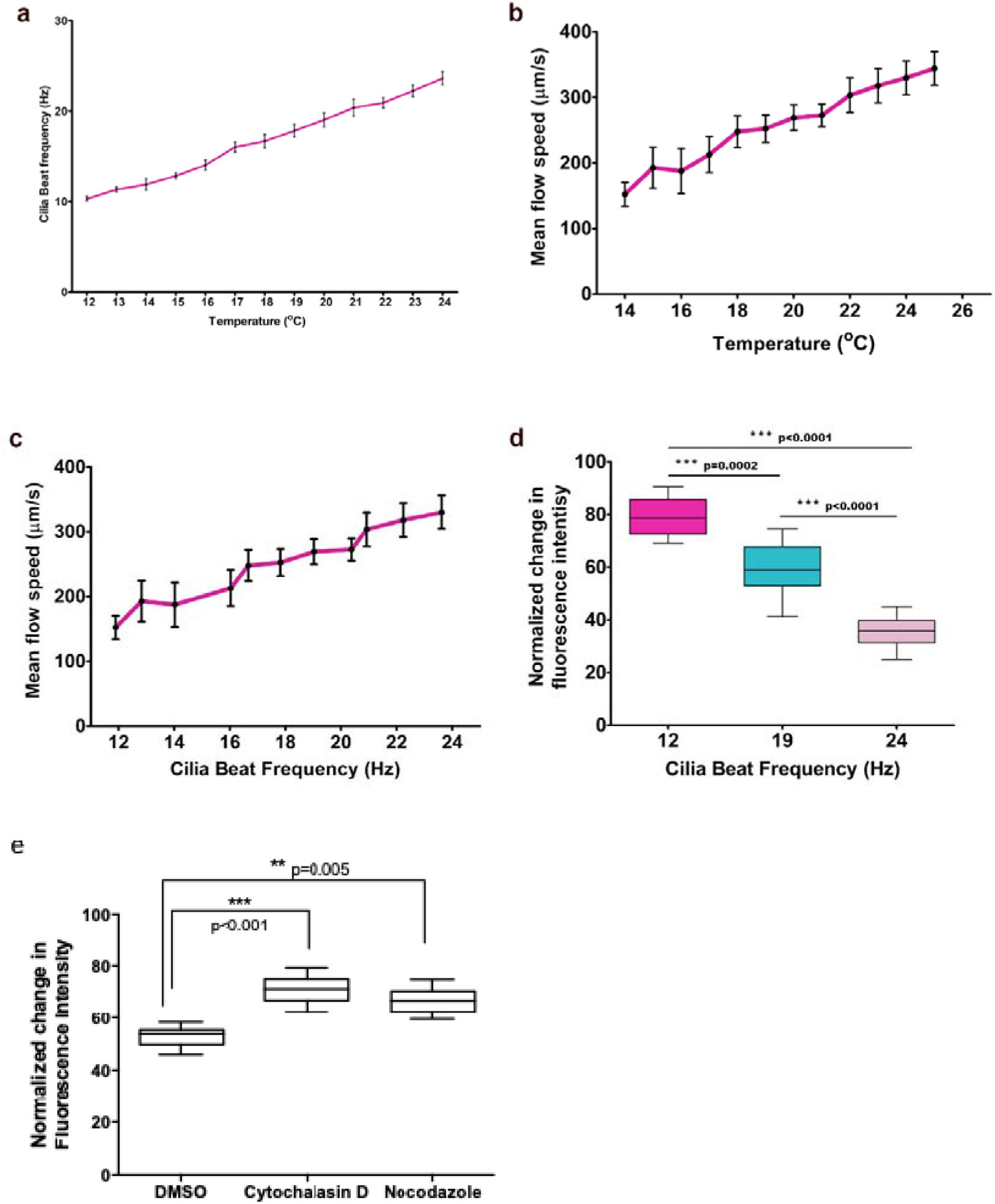
Detection of changes in flow attributed to cilia beat frequency or polarity modulation. a) Quantification of changes in *Xenopus* cilia CBF (n=5 tadpoles, 13 ROI from each tadpole), in response to temperature modulation. Temperature elevation induces increase in the frequency of *Xenopus* cilia beating. b) Quantification of changes in the flow generated by the mucociliary epidermis of *Xenopus* tadpoles (n=6 tadpoles), in response to temperature modulation. The velocity of flow is linearly correlated with temperature. c) Quantification of changes in flow velocity with respect to CBF modulation. The velocity of flow generated by the *Xenopus* mucociliary epidermis is proportional to CBF. d) End-point quantification of FITC intensity for tadpoles displaying differential CBF (n=6 tadpoles for each condition). Faster flow, resulting from CBF enhancement, accelerates the mixing of the biphasic system, as evident by more pronounced FITC quenching e) End-point quantification of FITC intensity for tadpoles treated with Cytochalasin D (n=5 tadpoles) or Nocodazole (n=5 tadpoles). Impaired flow, attributed to disturbed cilia orientation, results in slower mixing of the biphasic system, as evident by decreased fluorescence quenching.

As mentioned above the generation of robust flow depends on many factors including the rotational polarity, tissue level polarity, basal body spacing and beating pattern. The actin and microtubule networks have been shown to control these parameters and pharmacological disruption of either network using depolymerizing agents disrupts cilia polarity and negatively affects the generation of flow without affecting CBF (Werner et al., 2011).

To assess the capacity of the *Xenopus* flow assay to detect abnormal flow elicited by perturbed cilia polarity, we treated embryos with Cytochalasin D (10μM) or Nocodazole (1μM), that specifically target the sub-apical actin and microtubule network respectively, as previously described (Werner et al., 2011). Control and treated embryos were introduced in 96 well plates and end point flow assays were carried out. Treatment with either drug resulted in significant decrease in FITC quenching, resulting from defective flow generation, showing that the assay can detect defects stemming from deregulation of the cytoskeleton (**Figure 4e**).

Overall, these results show that the *Xenopus* flow assay can detect changes of flow velocity in a quantitative and linear fashion, but also changes influencing flow velocity in an indirect qualitative way (cilia polarity perturbation).

### The Xenopus flow assay can directly assess the effects of CBF-modulating compounds

The highly coordinated and directional beating of cilia, required for effective flow generation, is regulated by several factors, including calcium, ATP, nitric oxide and shear stress, acting via several signaling pathways (Salathe, 2007). To date, a large number of studies have identified compounds that act through these signaling pathways and either stimulate or inhibit CBF in human ciliated cells cultured in ALI systems (Workman and Cohen, 2014). However, due to the limitations of *in vitro* cell culture setups, which do not allow the direct assessment of changes in mucociliary flow, almost all of these studies focused strictly on monitoring changes in CBF. Although CBF is critical for the efficient generation of flow and should be therefore evaluated in studies of candidate ciliostimulatory compounds, the most relevant physiological parameter that should be assessed in such studies, is the flow generated by the mucociliary epithelium. Taking advantage of the *Xenopus*-based flow assay, we decided to test known ciliostimulatory compounds and directly assess their impact on mucociliary flow generation.

Following a systematic literature review we identified cAMP (Di Benedetto et al., 1991; Haxel et al., 2001; Runer and Lindberg, 1999; Schmid et al., 2006), IL-5 (Grosse-Onnebrink et al., 2016), IFN-γ (Grosse-Onnebrink et al., 2016), Prostaglandin-E2 (Bonin et al., 1992; Haxel et al., 2001; Schuil et al., 1995) and Sildenafil (Fowler et al., 2020), as compounds previously shown to enhance human ciliary motility either via changes in calcium levels or through various signal transduction pathways. *Xenopus* tadpoles (stage 34) were pre-treated with the candidate compounds, at concentrations indicated in **Figure 5**, and examined for their specific effects in mucociliary flow. Pre-treated tadpoles were introduced into individual wells of 96-well plates together with the low-density Biotin-4-FITC and high-density streptavidin solutions, both containing the respective compounds at the concentrations used for pre-treatment. FITC intensity in individual wells was measured every 5 minutes, for 1 hour. Quantification of fluorescence intensity over time, revealed that cAMP, IL-5, IFN-γ and Prostaglandin-E2 treatment accelerated quenching, resulting from solution mixing, indicating enhanced mucociliary flow (**Figure 5a**). However, Sildenafil treatment did not affect the mixing rate, compared to controls.

**Figure 5:**
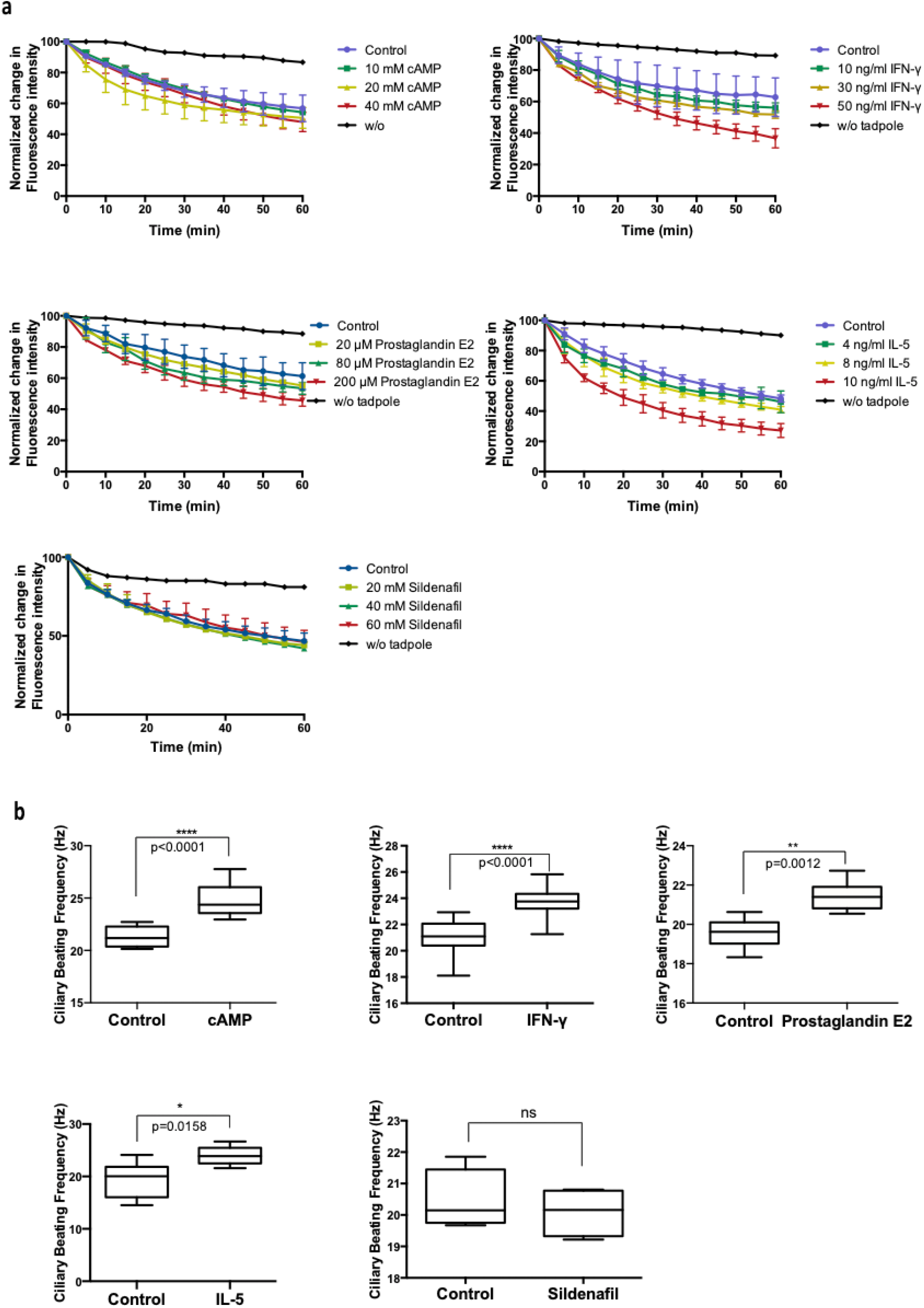
Treatment with cAMP, IL-5, IFN-γ and Prostaglandin-E2 enhances the beating frequency of Xenopus cilia and promotes faster mixing of solutions of differential density. *a) Xenopus* tadpoles treated with the indicated concentrations of cAMP (n=4), IL-5 (n=4), IFN-γ (n=4) and Prostagladin-E2 (n=4). Quantification of changes in FITC intensity show that treated tadpoles promote faster mixing of the FITC and streptavidin solutions, resulting in faster FITC quenching. Sildenafil treatment (n=4) did not lead to flow rate changes. b) *Xenopus* tadpoles treated with 20 mM cAMP, 10 ng/ml IL-5, 30 ng/ml IFN-γ, or 80 μM Prostaglandin-E2 display increased CBF compared to control, non-treated tadpoles. Treatment with Sildenafil (60 mM) did not affect CBF. (n=6 tadpoles for each condition)

To assess whether changes in mixing rate reflect CBF changes, treated tadpoles were imaged using brightfield high-speed video microscopy, followed by frequency analysis to determine the CBF under different conditions. In agreement with previous studies on mammalian cells, treatment with cAMP, IL-5, IFN-γ and Prostaglandin-E2 induced enhancement in CBF of *Xenopus* cilia, unlike Sildenafil which did not significantly change CBF (**Figure 5b**). In agreement with studies in mammalian cells, Prostaglandin-E2 induces ∼10% increase in CBF (Bonin et al., 1992) while IL-5 results in ∼20% (Grosse-Onnebrink et al., 2016). Using our assay, we observed ∼15% and 20% increase in mixing efficiency, respectively. However, treatment with IFN-γ results in ∼10% increase in CBF, followed by∼26% increase in mixing efficiency, compared to control tadpoles. This might be due to the regulatory role of IFN-γ in mucus production (Cohn et al 1999). On the other hand, CBF of Sildenafil-treated tadpoles remained unchanged compared to controls, in agreement with our *Xenopus* flow assay. Collectively, these data suggest that the *Xenopus* flow evaluation assay can quantitatively track the effects of CBF-modulating compounds.

## Discussion

Motile cilia lining the human airways are critical for mucociliary clearance leading to the removal of pathogens, cell debris and inhaled pollutants trapped in mucus, thus constituting a key component of the lung innate defense mechanisms (Ibanez-Tallon et al., 2003). Strong interest in cilia has emerged because of a growing number of diseases caused by defects in their formation and function. The best characterized disease regarding motile cilia is PCD, accompanied by recurrent respiratory infections and development of bronchiectasis, abnormally positioned internal organs, and infertility (Ibanez-Tallon et al., 2003; Noone et al., 2004).

As a rare disease, PCD lacks a specific treatment and many aspects of patient care are empirically based on therapeutic schemes applied for other chronic respiratory diseases, such as cystic fibrosis (Lucas et al., 2014). Such approaches rely on airway clearance therapies and aggressive antibiotic treatments (Joskova et al., 2020; Lucas et al., 2014). Some of the drugs used to improve airway clearance display secondary cilio-stimulatory properties, enhancing CBF, which is considered to further refine mucus clearance (Joskova et al., 2020). In fact, a lot of studies have identified compounds with similar properties, however, their direct impact on mucociliary flow has not been assessed, while their *in vivo validation* is limited (Joskova et al., 2020; Lucas et al., 2014; Workman and Cohen, 2014). PCD-specific drug development is challenging and therefore limited, because of the heterogeneity of the disease. At the same time, the lack of systems enabling direct assessment of mucociliary clearance in a fast, yet consistent way, complicates library screening for the identification of new compounds with positive impact on the effectiveness of mucociliary flow. Traditional methods for flow assessment require a combination of specialized, technically demanding and time-consuming approaches. Such methods, though, do not have the capacity to be applied for multiscale screenings to quantitatively validate the effects of candidate drugs on flow.

In this study, we developed a novel assay for the assessment of mucociliary flow, taking advantage of the external nature of the mucociliary epithelium found on *Xenopus* tadpoles’ skin. Unlike current methods, the *Xenopus* flow evaluation assay enables direct flow assessment in an easily reproducible, quick and cost-effective way, facilitating application in high-throughput screens for the identification of new therapeutic compounds for PCD.

The assay relies on the ability of *Xenopus* tadpoles to promote mixing of a system consisting of two solutions of differential density. This is visualized upon fluorescence labeling of one of the two solutions. Mixing, promoted by the robust flow generated by the tadpoles’ mucociliary epithelium, results in the diffusion of fluorescence, while the two phases initially generated by the two solutions are no longer visually distinguishable. In order to empower quantitative assessment of mixing efficiency, the Biotin-4-FITC/Streptavidin quenching system was introduced into this set-up. Specifically, Biotin-FITC (fluorophore) is introduced into the low-density solution, while Streptavidin (quencher) is introduced in the high-density solution. We show that mixing of the two solutions promotes the interaction between the fluorophore and the quencher, which is followed by measurable reduction of fluorescence intensity. As expected, fluorescence intensity changes are proportional to incubation time, indicating that they reflect changes in the extent of mixing. Addition of excess biotin (not influencing fluorescence in any way), results in termination of fluorophore-quencher interaction, enabling accurate and consistent end-point evaluation of the extent of mixing, thus facilitating the application of the assay in large scale experiments.

Our data show that the assay described in this study has the sensitivity to detect and quantify changes in flow velocity, resulting from defective mucociliary function. Specifically, we show that it can distinguish between normal and impaired flow, resulting from downregulation of FAK followed by defects in cilia density and function. In fact, differences in the severity of MCC phenotype are reflected by respective differences in the rate of mixing. This indicates that the rate of mixing is proportional to the rate of mucociliary flow and supports that the assay enables specific and reliable quantification of flow effectiveness. More importantly, our work demonstrates that quantification of flow-promoted mixing facilitates tracking of changes in parameters that influence the mucociliary flow, either quantitatively or qualitatively. In addition to proper cilia density, other parameters are critical for efficient flow generation, including CBF (influencing flow in a quantitative way), as well as CBP and cilia polarity (influencing flow in a qualitative way). In particular, we provide evidence that flow acceleration upon CBF enhancement is followed by measurable acceleration of solution mixing, showing direct correlation between CBF and mucociliary flow effectiveness. In addition, we show that the rate of mixing reflects flow defects, resulting from disruption of cilia polarity. This is a unique characteristic of the *Xenopus* flow evaluation assay, which therefore provides a valuable tool among current alternatives.

Finally, we validate the assay for its capacity to detect the effects of ciliostimulatory compounds on mucociliary epithelium. Such compounds have been shown to enhance CBF and are therefore considered to improve mucociliary clearance. However, the effect of such compounds on mucociliary flow at the tissue level, *in vivo*, cannot be easily assessed. Our work shows that the effects of CBF-modulating compounds are directly determined through the *Xenopus* flow evaluation assay. We show that CBF enhancement upon treatment with cAMP, IL-5, IFN-γ and Prostaglandin-E2 is directly correlated to enhanced mucociliary flow. Interestingly, we show that the magnitude of CBF increase is accompanied by relative acceleration of the biphasic solution mixing rate, suggesting that the assay provides an ideal tool for screening studies and the quantitative evaluation of the specific effects of ciliostimulatory compounds.

Overall, the methodology described in this paper provides a unique tool for the fast and straightforward quantification of mucociliary flow effectiveness. As such, it facilitates high-throughput screening of compounds as potential drugs for PCD, and at the same time the study of motile cilia function.

## Acknowledgements

We thank Professor Panayiotis Yiallouros (University of Cyprus) for kindly providing the Basler scA640 video camera and the Sisson-Ammons system, used for the analysis of cilia beat frequency.

