## Supplementary material for "Development of a *Xenopus*-based assay for high-throughput evaluation of mucociliary flow": (Supplementary figure 1)

**Supplementary Figure 1**

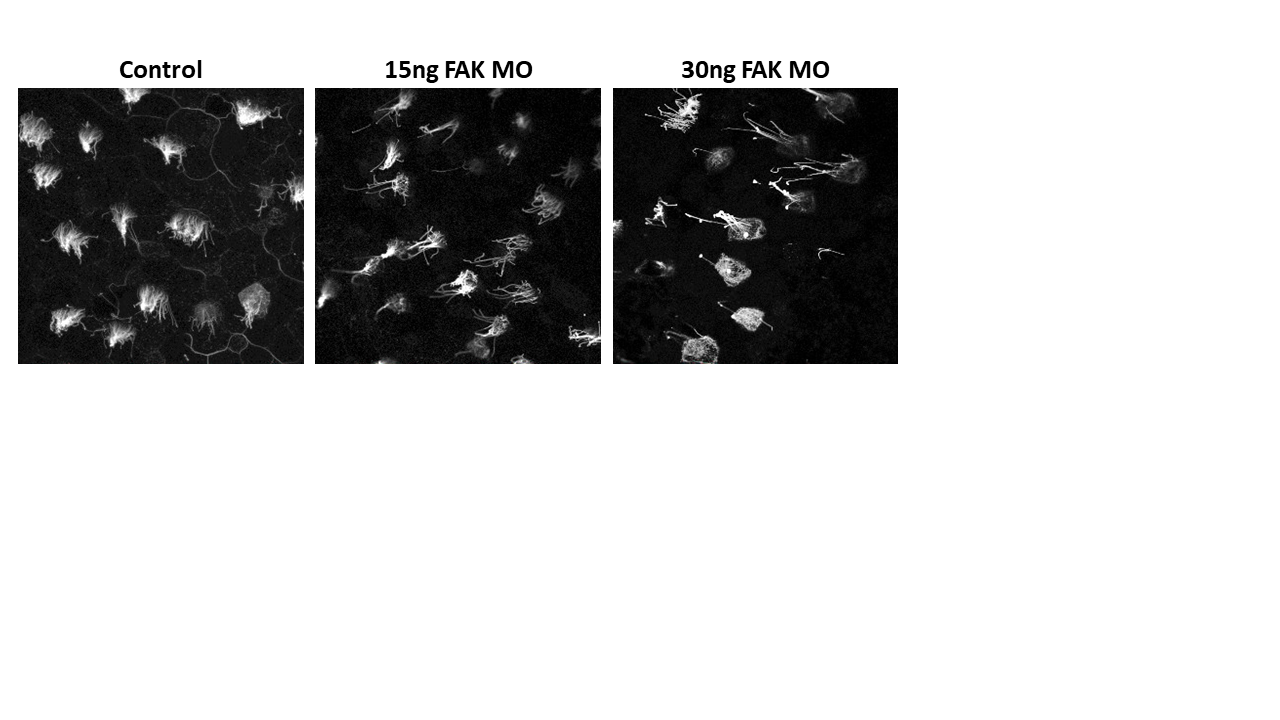

***Supplementary Figure 1: Downregulation of FAK elicits defects in multiciliated cell differentiation***

Representative maximum intensity projections of skin areas of control and FAK MO-injected tadpoles, stained for acetylated tubulin. Downregulation of FAK elicits dose dependent defects in multiciliated cells, which project less cilia compared to control cells.
